# *In vitro* cytokine expression analysis by droplet microfluidics

**DOI:** 10.1101/2021.12.28.474355

**Authors:** Ada Hang-Heng Wong, Semih Can Akincilar, Joelle Chua, Dhakshayini d/o K. Chanthira Morgan, Dorcas Hei, Vinay Tergaonkar

## Abstract

Droplet microfluidics provides a miniaturized platform to conduct biological assays. We previously developed a droplet microfluidic chip assay for screening cancer cells against chemical drugs and chimeric antigen receptor T (CAR-T) cells, respectively. In this study, we investigated chip application on a cytokine expression assay using MCF7 breast cancer reporter cells engineered by fusing green fluorescent protein (GFP) to the C-terminus of endogenous interleukin-6 (IL6) gene. Combined tumor necrosis factor α (TNFα) treatment and serum-free medium starvation stimulated IL6-GFP expression and enhanced GFP fluorescence. Our data showed that on-chip assay recapitulates the cellular response *in vitro*, although absolute quantification of IL6 induction could not be accomplished. The demonstration of multi-timepoint IL6 expression analysis paves the way for our future study on tumor response to immune attack via cytokine signaling.

## Introduction

In our early pursuit, we developed a droplet microfluidic assay to conduct drug screening on human primary cancer samples ^1^. Next, we tested if our chip can be applied to adoptive cell transfer applications ^2,3^ using anti-CD19 chimeric antigen receptor T (CAR-T) cells ^4^ as model. After establishment of the CAR-T assay, we noted the capability of our chip to interrogate a multitude of conditions, including multi-cell type analysis, variant cell-cell ratios, cell morphology analysis, *etc*. Hence, we went on to investigate the application of cytokine expression analysis to interrogate the feasibility of monitoring real-time protein expression on chip.

Cytokine signaling elicits a contact-free mechanism in the crosstalk between cancer cells and its microenvironment during cancer development and progression ^5–8^, which is suitable for analysis in cell suspension on our microfluidic chip. Interleukin 6 (IL6) is an important cytokine secreted by cancer cells to stimulate tumor proliferation by promoting tumor growth and creating a favorable tumor microenvironment *in vivo* in many cancer types, particularly breast cancer ^9–12^, which constitutes our primary research focus ^13,14^. Furthermore, IL6 is rapidly stimulated by tumor necrosis factor α (TNFα) in minutes and its expression sustains for hours after stimulation in certain cell lines, thus provides the necessary time window for data collection on chip. Subsequently, digital screening of IL6 expression on public data ^15^ and quantitative PCR (qPCR) analysis led to the choice of the luminal breast cancer cell line MCF7. However, MCF7 belongs to epithelial cell type, which are often grown in adherent cultures or embedded in biogels, e.g. Matrigel^®^. It is arguable that MCF7 cells may not display the necessary immune response in suspension. Nevertheless, the fact that MCF7 cells form mammospheres ^16^, an important assay to demonstrate tumorigenicity in breast cancer research, suggests that this cell line is adaptable to suspension culture. Most importantly, MCF7 cells can be specifically targeted by CAR-T cells ^17^. Altogether, these properties make MCF7 cells a desirable model for our research.

Hence, we constructed an MCF7 reporter cell line that expresses IL6 endogenously fused with green fluorescent protein (GFP) to conduct a multi-timepoint, on-chip cytokine expression analysis *in vitro* (Figure 1a).

**Figure 1.**
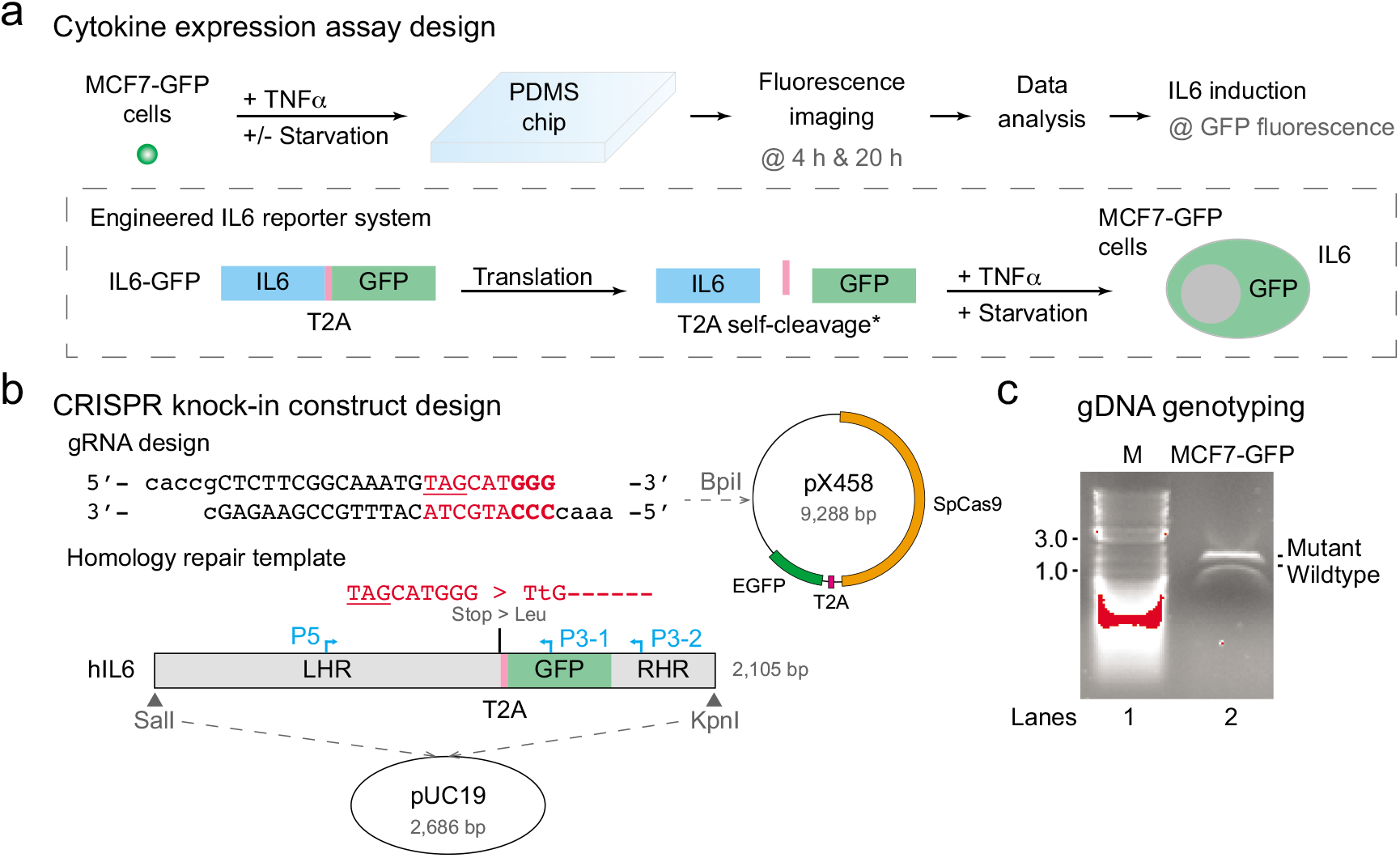
IL6-GFP reporter assay. (**a**) Overview of a cytokine expression assay design. MCF7 breast cancer cells were engineered to express a fusion protein of IL6-GFP (inset) after stimulation by TNFα treatment and serum-free medium starvation. GFP fluorescence was captured by fluorescence imaging for assessment of reporter activation. (**b**) Construct design for the IL6-GFP knock-in plasmids by CRISPR technology. The guide RNA (gRNA) for SpCas9 targeting was designed against the C-terminus of human *IL6* locus (Ensembl GRCh38) and inserted into pX458 vector after vector digestion by BpiI. The homology repair template was designed by cloning the T2A peptide (T2A) and GFP between the 1,000 bp left homology arm (LHR) and the 316 bp right homology arm (RHR) designed in-frame against the two sides of the stop codon of human *IL6* gene, followed by insertion into the pUC19 vector after double digestion by SalI and KpnI. Please note that the repair template consisted of a single nucleotide mutation of TAG > TtG to remove the stop codon of human *IL6* gene and a 6-nt deletion to remove the protospacer adjacent motif (PAM) recognized by SpCas9. Mutated sequences are highlighted in red; the stop codon of human *IL6* gene is underlined; the PAM is highlighted in bold text; the translated sequence of the repair template is indicated in grey; the primers used for PCR genotyping is shown in cyan. (**c**) PCR genotyping of MCF7-GFP cells. PCR genotyping was conducted on genomic DNA (gDNA) extracted from MCF7-GFP cells using P5 and P3-2 primers. Both the mutant and wildtype bands were gel extracted and sequence validated by Sanger sequencing to confirm the heterozygous insertion of GFP in *IL6* locus.

## Materials and Methods

### Construction of MCF7-GFP reporter cell line expressing IL6-GFP

MCF7-GFP cells were constructed by CRISPR knock-in via homologous recombination (Figure 1b). First, GFP was fused to the C-terminus of endogenous *IL6* gene to enable transcriptional activation of the endogenous IL6 promoter by nuclear factor kappa B (NFκB) signaling ^11^ at 1:1 IL6:GFP expression. Second, C-terminal GFP insertion was chosen to preserve the N-terminal secretory peptide, so that the normal regulation of IL6 secretion might not be compromised. Third, addition of the T2A peptide ^18^ was intended to separate the IL6 and GFP proteins to reduce protein size to minimize the impact of GFP fusion on the extracellular secretion of IL6. Moreover, our hypothesis was that GFP would be retained in cytosol for intracellular fluorescence detection on chip. Fourth, the stop codon was removed to enable GFP expression and, together with the deletion of the adjacent nucleotides, minimizes the chance of SpCas9 cleavage to increase recombinant efficiency. Finally, the fluorescence of GFP protein was used to detect cytokine induction on chip.

Briefly, MCF7 cells were co-transfected with the pX458 vector (Addgene #48138) co-expressing gRNA, SpCas9 and EGFP, and the pUC19 vector (New England BioLabs #N3041) containing the repair template. Transfected cells transiently expressing EGFP of pX458 vector were sorted out by fluorescence activated cell sorting (FACS) and genotyped by polymerase chain reaction (PCR). Sanger sequencing of the two DNA bands amplified from genomic DNA (Figure 1c) was performed to validate the genomic sequence of the positive clone.

### Droplet microfluidic chip assay

The droplet microfluidic chips were fabricated from polydimethylsiloxane (PDMS) by soft photolithography and loaded as previously described ^1^. Briefly, each chip was fabricated to contain two channels of 48 wells where samples were pre-mixed and loaded into the inlet. Water-in-oil droplets form in each well due to fluid exclusion at the channel restriction. Thus, one large droplet (25-35 nL) containing the sample cells form in each well of the chip, yielding a maximum of 48 experimental replicates for each channel. The cells were incubated in suspension within the droplets. Loaded chips were incubated in a 37°C humidified incubator supplemented with 5% CO_2_. Fluorescence imaging was performed using 10× objective on Carl Zeiss LSM 800 Confocal Microscope. All images were processed by ImageJ v1.52e in TIFF format.

### Time-lapse live cell imaging

1.0 × 10^5^ MCF7-GFP cells were seeded on a 35 mm confocal dish. After the cells adhered, they were starved in OptiMEM™ medium for 12 h. Hoechst staining was performed in Dulbecco’s phosphate buffer saline (DPBS) at room temperature for 30 min before replacing the medium with 1mL OptiMEM™ medium supplemented with 10 pg/mL TNFα. Next, the dish was equilibrated on stage in the 37°C humidified chamber supplemented with 5% CO_2_ of Carl Zeiss LSM 800 Confocal Microscope for 30 min. Time-lapse live cell imaging was performed under 10× DIC, DAPI and GFP channels at 30 min intervals for 8 h duration. Raw images were saved as CZI and processed by ImageJ v1.52e.

### mRNA and protein quantification

For mRNA quantification, 1.0 × 10^5^ MCF7-GFP cells were seeded in each well of a 12-well plate and cultured overnight in DMEM medium before starvation for 12 h in OptiMEM™ medium. Starved MCF7-GFP cells were treated with 10 pg/mL TNFα in OptiMEM™ medium for the time as indicated in figure legends. Adherent cells were collected after trypsinization and total RNA was extracted by TRIzol™ (Life Technologies) and purified by affinity column purification (Macherey-Nagel). Reverse transcription was performed using Maxima Reverse Transcriptase (Thermo Fisher). qPCR was performed on Bio-Rad CFX96 Touch Real-Time PCR Detection System using SYBR^®^ Green Master Mix (Bio-Rad) following manufacturer’s protocol.

For protein quantification, 4.0 × 10^5^ MCF7-GFP cells were seeded in each well of a 6-well plate and cultured overnight in DMEM medium before starvation for 12 h in OptiMEM™ medium. Starved MCF7-GFP cells were treated with 10 pg/mL TNFα in OptiMEM™ medium for the time as indicated in figure legends. Culture medium supernatant was collected and subjected to enzyme-linked immunosorbent assay (ELISA) of secreted IL6 (R&D Systems) detected by absorbance at 450 nm and 570 nm on Molecular Devices SpectraMax M2 Microplate Reader. For Western blot, secreted proteins were enriched by chloroform-methanol extraction or by immunoprecipitation by anti-IL6 antibody (data not shown) from culture medium supernatant, whereas intracellular proteins were collected from adherent cells lysed in SDS loading dye upon heating at 95°C. Western blot was performed on polyvinylidene fluoride (PVDF) membranes using anti-human IL6 antibody (Santa Cruz #sc-7930), anti-GFP antibody (Abcam #ab290) or anti-β-actin antibody (Sigma-Aldrich #A2066), and detected by chemiluminescence using Clarity™ Western ECL Substrate (Bio-Rad) on Bio-Rad ChemiDoc™ Touch Imaging System.

### Flow cytometry analysis

5.0 × 10^5^ MCF7-GFP cells were seeded in each well of a 6-well plate and cultured overnight in DMEM medium before starvation for 12 h in OptiMEM™ medium. Starved MCF7-GFP cells were treated with 10 pg/mL TNFα in OptiMEM™ medium for 4 h, trypsinized and subjected to FACS on BD LSRFortessa™ Flow Cytometer. Results were exported as FCS files and analyzed by Flowing Software v2.5.1.

### Data analysis and visualization

All graphs and plots were drawn by GraphPad Prism v5.1 or using *ggplot* in R v3.3.2. Figures were prepared by assembling images, graphs and plots using Adobe^®^ Illustrator^®^ CS6 v16.0.0.

## Results

In this study, we applied TNFα and serum-free medium starvation to stimulate IL6-GFP protein expression in MCF7-GFP reporter cells (Figures 1a & 1b). IL6-GFP reporter activation was interpreted by calculating the ratio of GFP^+^ cells to all cells in each droplet on chip. Results showed that the mean percentage of GFP^+^ cells at 4 h post-treatment was significantly higher in MCF7-GFP cells treated with TNFα and starvation as compared to TNFα-treated cells without starvation, whereas insignificant difference between the two conditions were observed at 20 h post-TNFα treatment (Figure 2a). This result was consistent with qPCR analysis, which showed that both *IL6* and *GFP* mRNA expression surged at 1 h post-TNFα treatment (Figure S1a). Notably, the relative induced mRNA level of *GFP* was half of that of *IL6*, in concordance to the heterozygous *GFP* insertion in MCF7-GFP cells verified by PCR genotyping (Figure 1c). On the protein level, Western blot analysis of protein expression showed that there was a leak expression of IL6-GFP under untreated conditions, whose expression increased after starvation and remained level after TNFα treatment in the intracellular fraction (Figure S1b). The observed band size suggested that the fusion protein was expressed, but it was not cleaved at the T2A peptide as designed. No band specific to our construct was detected in culture medium on Western blot (Figure S1b). Endogenous IL6 protein was neither detected in cell pellet nor culture medium on Western blot (Figure S1b). ELISA performed on the culture medium supernatant of MCF7-GFP cells after TNFα treatment and starvation showed that secreted IL6 was detectable only after 4 h post-TNFα treatment and increased to 1.3 pg/mL at 8 h post-TNFα treatment (Figure S1c). However, it remained inconclusive whether ELISA detected the endogenous IL6 or IL6-GFP protein, provided that neither of them was detected in cell culture medium supernatant on Western blot.

**Figure 2.**
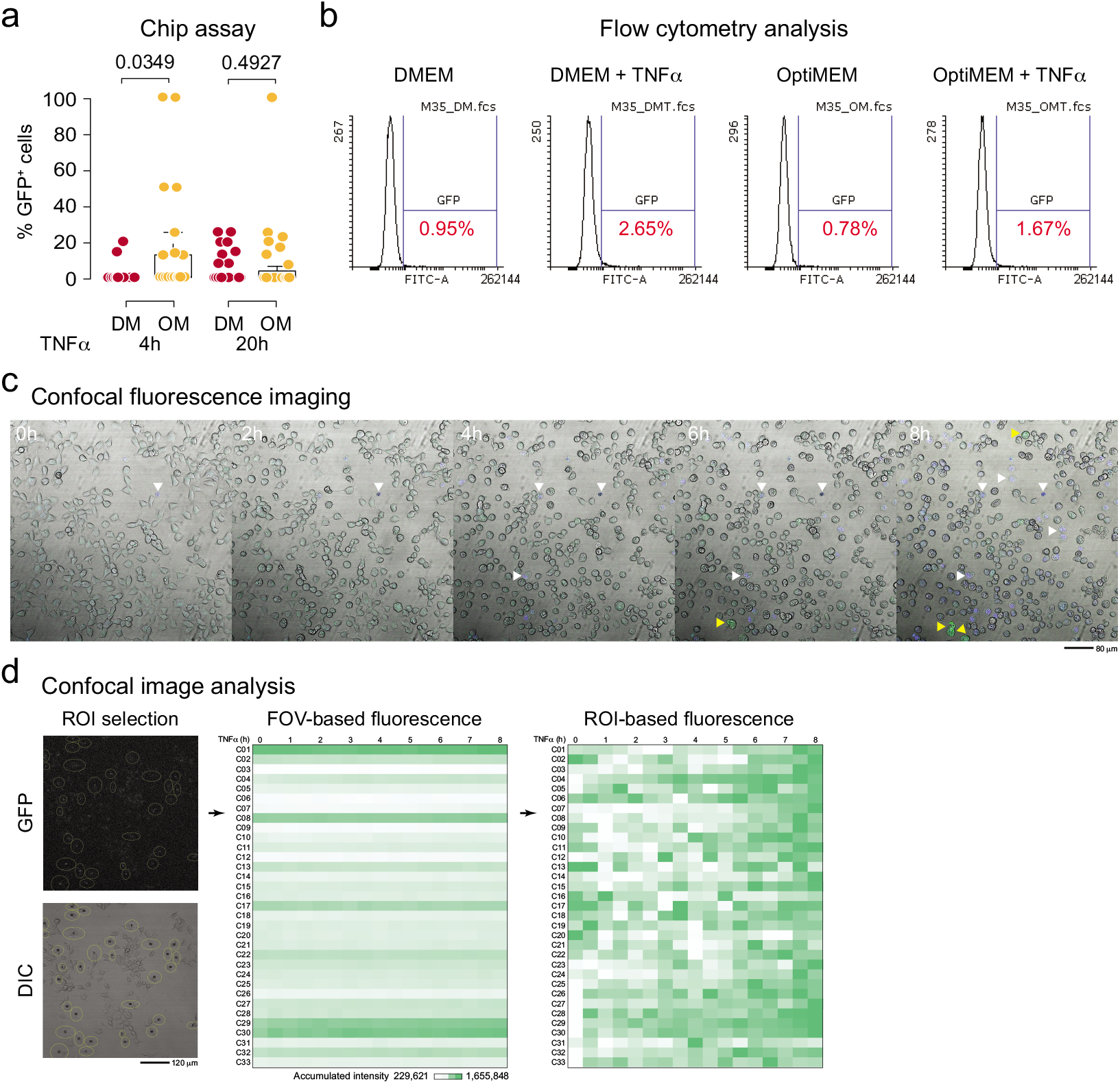
Analysis of IL6-GFP reporter system. (**a**) Reporter activation assay on chip. MCF7-GFP cells were loaded on chip and imaged at 4 h and 20 h post-TNFα treatment in full culture medium DMEM (DM) or serum-free OptiMEM™ medium (OM). Each dot represents one microfluidic well, the thick black line, grey box and dotted error bars of the box plot indicate median droplet cell count, interquartile range between −1σ and 1σ, and range of droplet cell count of all wells on chip, respectively; the numbers indicate *p* values of 1-tailed *t* test. (**b**) Flow cytometry analysis of MCF7-GFP cells. MCF7-GFP cells were trypsinized and subjected to FACS analysis after stimulation by serum-free medium starvation with or without TNFα treatment. The percentage of GFP^+^ cells calculated by identical cutoff across all samples is shown. (**c**) Time-lapse live cell imaging of MCF7-GFP cells after stimulation by TNFα treatment and serum-free medium starvation. Confocal fluorescence imaging was performed under 10× DIC, DAPI and GFP channels at 30 min intervals for 8 h duration. White arrows indicate Hoechst-stained cells, indicative of cell death; yellow arrows indicate GFP^+^ cells, indicative of reporter activation. (**d**) Analysis of GFP fluorescence of MCF7-GFP cells by time-lapse live cell imaging on confocal microscope. MCF7-GFP cells were imaged under 20× DIC and GFP channels on confocal microscope at 30 min intervals up to 8 h post-TNFα treatment. Single cells were randomly selected for fluorescence quantification. Accumulated fluorescence intensity was calculated by multiplying mean GFP fluorescence intensity by ROI area, the absolute values of which is shown under the FOV-based fluorescence heatmap. Please note that the absolute accumulated intensity of FOV-based fluorescence was scaled according to the FOV, whereas the relative accumulated intensity of ROI-based fluorescence was scaled according to each ROI manually segmented as a single cell. The GFP and DIC images indicate the position of each ROI used for fluorescence intensity analysis; the rows and columns of the heatmap indicate the ROI and acquisition time post-TNFα treatment, respectively; the scale bar indicates absolute accumulated fluorescence intensity in FOV-based fluorescence image.

To compare our chip assay with conventional methods, flow cytometry analysis and time-lapse fluorescence imaging were used. Flow cytometry analysis is a robust, high-throughput method for reporter analysis, but it does not support real-time monitoring nor morphological analysis. Alternatively, time-lapse fluorescence imaging provides a better resolution on both the spatial and temporal scale, but it lacks throughput as compared to flow cytometry. Consequently, flow cytometry analysis showed insignificant difference between TNFα-treated and untreated cells in both unstarved and starved conditions (Figure 2b). On the other hand, time-lapse live cell imaging showed that GFP fluorescence increased in some MCF7-GFP reporter cells that did not undergo cell death (Figure 2c). Fluorescence quantification of randomly selected single cells at different locations in the field of view showed that each cell displayed its own fluorescence intensity and fluorescence fluctuation pattern over time as compared to the other cells (Figure 2d). Nevertheless, GFP fluorescence of the analyzed cell population depicted an overall increasing trend up to 8 h post-TNFα treatment, partially due to rounding up of cells under stress yielding more voxels on the z-axis for fluorescence signal detection. On this aspect, our chip assay eliminates the fluorescence signal fluctuation due to morphological change because cells are generally globular in suspension. Notably, the majority of MCF7-GFP cells died at 20 h post-TNFα treatment and starvation under the confocal microscope during time-lapse imaging (unpublished data), possibly because of cellular stress caused by TNFα treatment, starvation and phototoxicity. Taken together, we deduced that the lack of synchronized and sharp GFP fluorescence induction in the MCF7-GFP cells after TNFα treatment and starvation led to failure of applying the reporter system on FACS. Hence, on-chip fluorescence analysis opens a new realm to analysis of reporters displaying subtle changes. This feature is reflected in the power of analysis of our chip assay because, in concordance to other single cell technologies, each cell is considered as one probabilistic event. Hence, filling one channel of 48 wells will conclude if there are any medium-to-weak effects (Figure S2).

## Discussion

The IL6 reporter assay demonstrated in this study shows that cytokine expression analysis is feasible on our microfluidic chip. This adds to our toolbox to study drug-cell, cell-cell and stimulus-response interactions that are fundamental to biological studies (Figure 3). However, there are certain technical limitations: (1) the current chip design does not support perfusion or addition of reagents after chip loading; (2) data acquisition is limited to 4 h intervals, which restricts its application on transiently expressed proteins, such as TNFα; (3) the application of fluorescence microscopy limits the number of reporters to the number of available fluorophores; (4) the long half-life of GFP (26 h in cells ^19^) indicates the induction of protein expression but not degradation, resulting in qualitative assessment of IL6 expression. Nevertheless, this assay enables quantitative assessment of how many cells displayed IL6 induction upon TNFα treatment and serum-free medium starvation at specific time points, which is not resolved by flow cytometry and conventional time-lapse fluorescence imaging.

**Figure 3.**
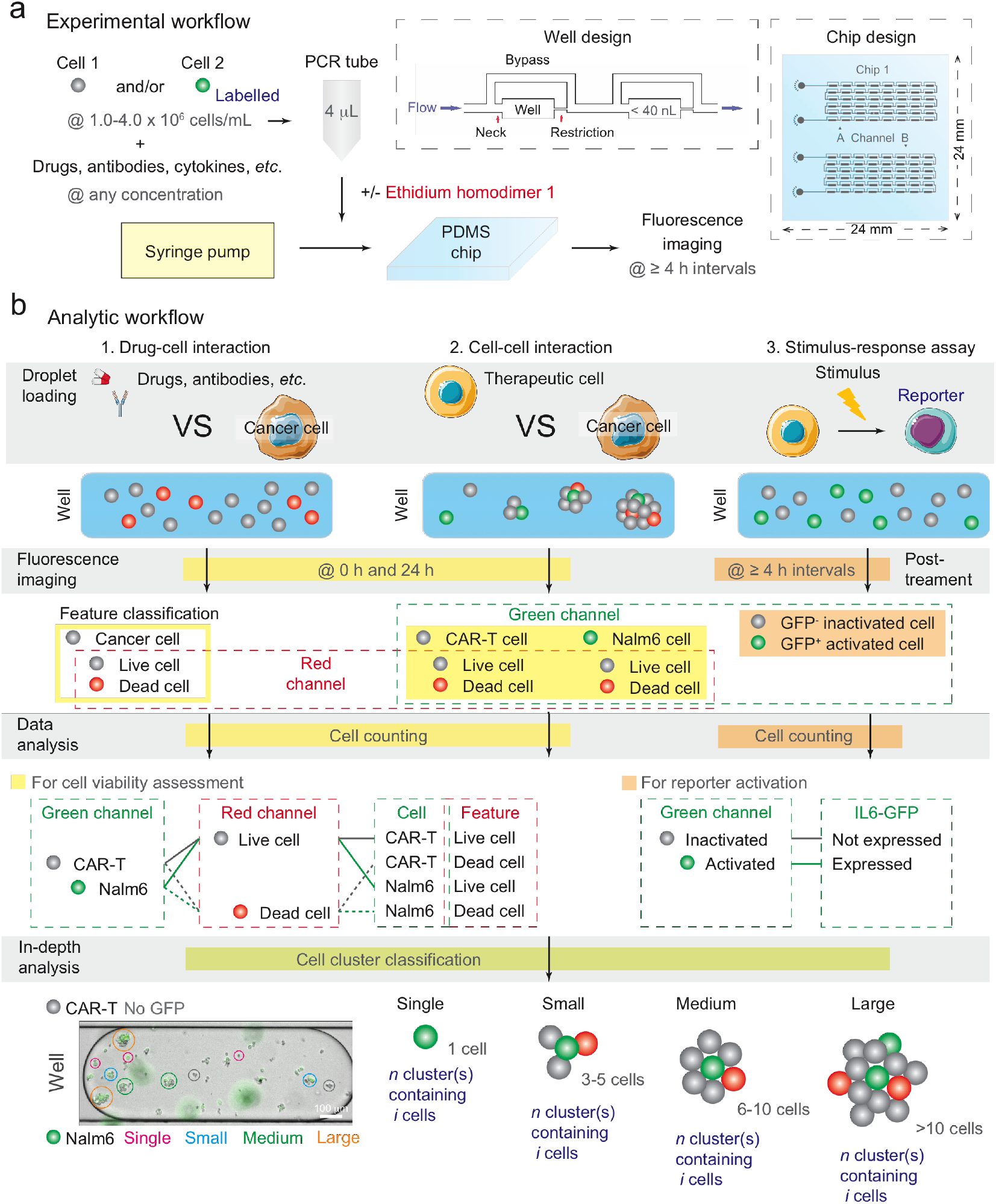
Overview of the droplet microfluidic chip assay. **(a)** Outline of experimental workflow. Cells, drugs, antibodies, cytokines, *etc*, are first mixed in a PCR tube and then loaded into one channel of a PDMS chip containing 48 microwells (inset), followed by fluorescence imaging at desired time points. Ethidium homodimer 1 may be added to indicate dead cells. **(b)** Outline of analytic workflow. After loading the necessary components on chip, fluorescence imaging under relevant fluorescence channels is conducted to acquire essential cell features for data analysis. For example, the green and red channels are used for CAR-T/Nalm6 and live/dead cell identification, respectively, in the CAR-T cytotoxicity assay, whereas the green channel is sufficient for assessing IL6-GFP reporter activation. Besides fluorescence, brightfield (BF), phase contrast (Ph) or differential interference contrast (DIC) images are acquired in all assays for cell counting purpose. In-depth analysis to study CAR-T cytotoxic mode was conducted by classifying aggregates of cells into different cell cluster types based on cluster size and/or cell count in these aggregates (BF image with annotations); other morphological classification methods can be used based on research needs.

IL6 is a pleiotropic cytokine, which shows pro- and anti-cancer effects depending on the cell model and culture conditions. For instance, elevated IL6 expression in MCF7 cells induces an epithelial-to-mesenchymal (EMT) phenotype ^20^, which is considered an indicator of tumor metastasis. Concordantly, IL6 secreted by cancer-associated fibroblasts is reported to trigger tamoxifen resistance in MCF7 cells ^21^. These data suggest that IL6 is pro-tumorigenic ^5^. Paradoxically, IL6 has pro-inflammatory roles to stimulate T cell activity ^22–25^. Hence, it remains elusive whether it is the secondary effects of tumor-secreting IL6, such as prevention of lymphocyte infiltration ^26^, disruption of T cell activation by other immune cells ^27,28^, or transformation of CD8^+^ T cells into a non-cytotoxic form ^29^, that contributes to the pro-tumorigenic action of IL6 observed in MCF7 cells.

IL6 elevation is a phenomenon of immune evasion, which often contributes to poor treatment outcome ^30^. Thus, understanding the biological mechanisms of IL6 elevation and potentially being able to perform drug screens to alleviate the condition becomes valuable. For this purpose, the high specificity, robustness and controllable plasticity of CAR-T cells provides an excellent tool to mimick aggressive and lethargic immune invasion. Therefore, our future goal is to apply the MCF7-GFP cells described here, together with other reporter cell lines, in the presence of CAR-T cells with different specificities on our microfluidic chip to dissect IL6 signaling in a clean *in vitro* system. Results may suggest whether IL6 would still be expressed by cancer cells in the presence of active or inactive CAR-T cells. Moreover, although IL6 usually induces a positive feedback loop in the tumor microenvironment ^31^, MCF7 cells display a relatively low basal and induced level of IL6 expression, suggesting that MCF7 cells possess some mechanism to repress IL6 expression ^11^. Hence, understanding the mechanism will enable us to overcome immune evasion reflected in IL6 elevation in MCF7 cells.

## Conclusion

In this study, we constructed an MCF7-GFP reporter cell line to conduct a multi-timepoint cytokine expression analysis on chip *in vitro*. It paves the way for our immune-tumor interaction studies in the future.

## Acknowledgements

Acknowledgements to Prof. Yanwei Jia of University of Macau for provision of the chip design and microfabrication facility.

## Author contributions

This project was conceived by A.H.W.. V.T. provided wet lab facilities. A.H.W. conducted all experiments and analyzed all data; J.C. provided bioinformatics support for digital screening of IL6-expressing cell lines; S.C.A. and D.C.M. provided technical assistance for CRISPR knock-in and FACS, respectively; D.C.M. and D.H. conducted the IP experiment. All authors prepared and approved this manuscript.

## Conflict of interest

A.H.W. received incubation funds from the University of Macau Development Foundation to setup the startup company AW Medical Company Limited.

## Data and Materials Availability

All data is available upon request from the corresponding authors.

## Supplementary Figures

**Figure S1.**
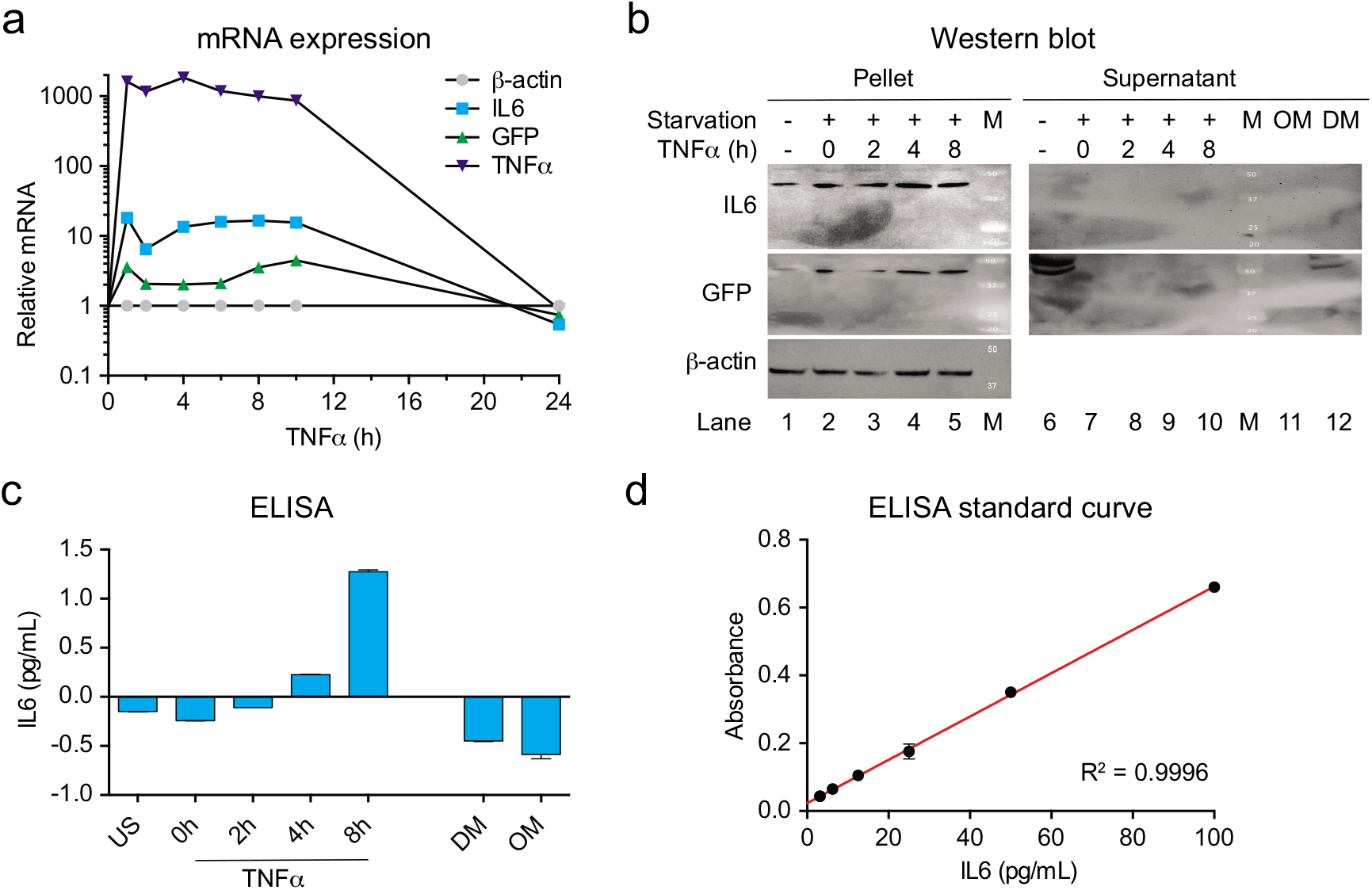
mRNA and protein quantification of IL6-GFP. (**a**) qPCR analysis of IL6-GFP induction after stimulation by TNFα treatment and serum-free medium starvation. Total RNA was extracted from MCF7-GFP cells after stimulation by TNFα treatment and serum-free medium starvation. IL6 and GFP mRNA surged at 1 h post-treatment. TNFα mRNA served as positive control; β-actin served as reference control. Each dot represents mean relative mRNA expression of triplicate wells; error bars indicate standard deviation of the mean. (**b**) Western blot analysis of IL6-GFP induction after stimulation by TNFα treatment and serum-free medium starvation. Intracellular (Pellet) and culture medium supernatant (Supernatant) fractions were collected from MCF7-GFP cells after 12 h starvation and indicated time points post-TNFα treatment. For IL6-GFP expression, Western blot was performed on a single membrane by blotting IL6 and GFP consecutively after stripping. M indicates protein marker, and corresponding molecular weight in kDa are indicated on the blot; OM and DM indicate sampling from OptiMEM™ and DMEM without cells, respectively. (**c**) ELISA of secreted IL6 after stimulation by TNFα treatment and serum-free medium starvation. Cell culture medium supernatant was collected from MCF7-GFP cells after stimulation by TNFα treatment and serum-free medium starvation, and subjected to ELISA. The bars indicate mean IL6 protein levels of duplicate wells; the error bars indicate standard deviation of the mean. (**d**) Standard curve for IL6 quantification in ELISA experiment. Serially diluted solutions of IL6 standard was assayed in duplicate wells in parallel to the ELISA experiment to derive the standard curve for IL6 quantification. Each dot represents the mean difference between the absolute absorbance of 450 nm and 570 nm of duplicate wells; the error bars indicate standard deviation of the mean; the red line of the standard curve indicates linear regression of the data points by GraphPad Prism.

**Figure S2.**
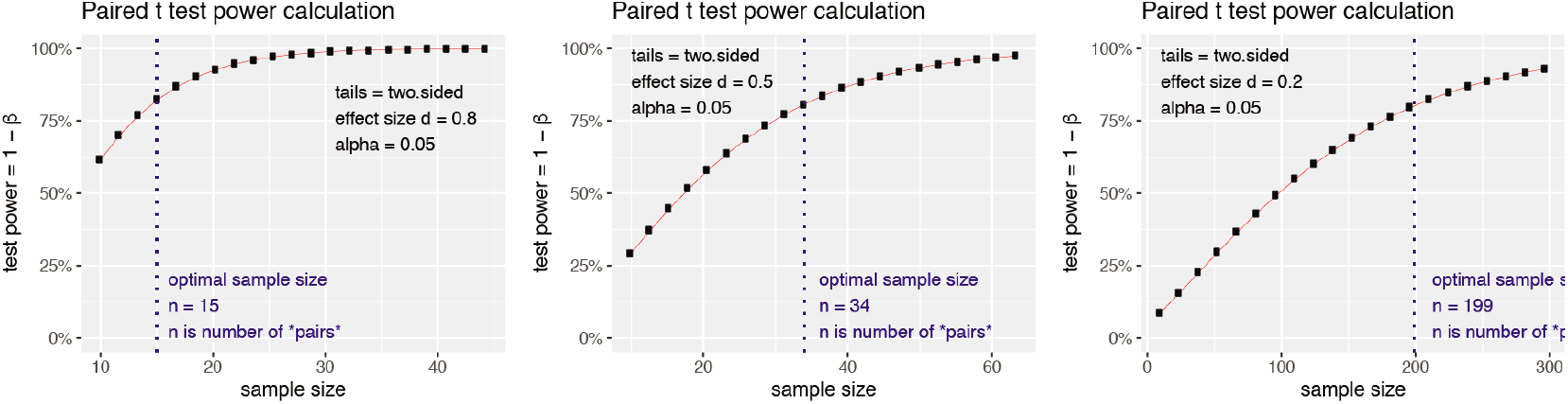
Power of analysis. Power of analysis was conducted using the R package *pwr* to calculate sample size under the following criteria: (a) each cell is considered as one probabilistic event; (b) α=0.05, i.e. applying the 95% confidence interval; (c) β=0.20, i.e. using 80% power of analysis; and (d) applying the generic effect sizes of d=0.8, 0.5 and 0.2 for large, medium and small effects, respectively. Results confirmed that our chip assay provides the power of analysis even for weak reporters.

